# Anti-mutagenic and synergistic cytotoxic effect of cisplatin and Honey Bee venom on 4T1 invasive mammary carcinoma cell line

**DOI:** 10.1101/168542

**Authors:** Faranak Shiassi Arani, Latifeh Karimzadeh, Seyed Mohammad Ghafoori, Mohammad Nabiuni

**Affiliations:** Department of Animal Biology, Faculty of Biological Science, Kharazmi University, Tehran, Iran; Laboratories’ Animal Center & Cellular and Molecular Research Laboratory,Faculty of Biological Sciences, Kharazmi University, Tehran, Iran; Department of Genetics, Islamic Azad University, Tehran medical Branch, Tehran, Iran; Department of Cell and Molecular Biology, Faculty of Biological Sciences, Kharazmi University, Tehran, Iran

**Keywords:** Ames assay, Cancer, cell culture, Cisplatin, MTT assay

## Abstract

Honey Bee Venom has various biological activities such as inhibitory effect on several types of cancer. Cisplatin is an old and potent drug to treat the most of cancer. Our aims in this study were determination of the anti-mutagenic and cytotoxic effects of HBV on mammary carcinoma, lonely and in combination with cisplatin. In this study 4T1 cell line were cultured and incubated at 37°C in humidified CO2-incubator. The cell viabilities were examined by MTT assay. Also HBV was screened for its anti-mutagenic activity against sodium azide by Ames test. The result showed that 6μg/ml HBV, 20μg/ml cisplatin and 6μg/ml HBV with 10μg/ml cisplatin can induce an approximately 50% 4T1 cell death. 7mg/ml HBV with the inhibition of 62.76% sodium azide showed high potential in decreasing the mutagenic agents. MTT assay demonstrated that HBV and cisplatin can cause cell death in a dose-dependent manner. The cytotoxic effect of cisplatin is also promoted by HBV. Ames test results indicated that HBV can inhibit sodium azide as a mutagenic agent. Anti-mutagenic activity of HBV was increased significantly in presence of S9 mix. Hence, our findings reveal that HBV can enhance the cytotoxic effect of cisplatin drug and it has cancer preventing effects.

## Introduction

Breast cancer is originated from the breast tissue and uncontrolled growth of cells from the inner lining of milk ducts or lobules^1^. Breast cancer is the most common diagnosed cancer and the main cause of cancer death among females worldwide^2^. Several studies have shown that breast carcinoma is occurred mostly in undifferentiated structures of breast and these structures are the origin site of ductal carcinoma. Approximately 1–2 years after onset of the first menstrual period, lobule formation is started. There is a gradual process for mammary gland which it needs several years. Parous women particularly women who have full term pregnancy experience at young age, have full lobular differentiation in breast structure ^3^ The most risk factors for breast cancer have been estimated and studied included diet, oral contraception, postmenopausal substituent treatment with estrogen, breast irradiation, and environment ^4^.

The 4T1 mouse mammary tumor cell line is a model of breast cancer which able to metastasize to site affected in human breast cancer. In 1983, Fred Miller and coworkers isolated the 4T1 cell line from BALB/c mammary tumor. This cell line has capacity to metastasize to bone and several organs affected in breast cancer consist of lungs, liver, brain. Hence in the recent years, usage of this cell line has increased^5^.

The molecular studies of mammary tumors indicated that the amplification of oncogenic genes such as *erb-B1, erb-B2, c-myc, int-2* are common ^6^ About 5–10% of all breast cancer cases are hereditary breast cancer. *BRCA1*, tumor suppressor gene, is mutated in the most hereditary breast and ovarian cancer ^7^

Recently treatment of cancer is a global concern. The various types of cancer treatment ways such as surgery, anticancer drug (chemotherapy), irradiation, hormone therapy, nutritional supplementation are used. Chemotherapy is a systematic therapy in which all of body cells are exposed to chemotropic agents^8^.

Usage of metals as medicine traces back to 5000 years ago. The studies on inorganic chemist develop modern medicine with metal components to treat disease such as cancer Cisplatin [cis-dichlorodiammineplatinum(II)] is a first stage of chemotherapy for most cancer including testicular, ovarian, cervical, small lung-cell and also breast cancer ^9^.

Nevertheless cytotoxic effect of cisplatin depends on dosage, and its high dosage has improvement effects on cancerous cells. However, usage of cisplatin is limited because it has several side effects specially nephrotoxicity and neurotoxicity effects^10^.

Honey bee venom (HBV) is an active product which is produced by the venom glands associated with the sting apparatus of honey bee workers and their queen. The history of apitherapy traces back 3000 to 5000 years ago ^11^. HBV is a very complex mixture of active enzymes, peptides and amines. Its most important components are Melittin and phospholipase A_2_, Adolapin and mast cell degranulating peptide. The several *in-vitro* and *in- vivo* studies revealed that HBV has antiinflammatory, cytotoxic and antibacterial effects and also it can cause severe allergic reaction ^12^. HBV as an old medicine has been used to treat arthritis, rheumatism, back pain, cancerous tumors, and skin diseases ^13^.

Our aims in this study were attended to investigate the cytotoxic effect of honeybee venom on 4T1 cell line, lonely and in combination with cisplatin, and also analysis of anti-mutagenic activity and anti-cancerous effects of bee venom by Ames test.

## Materials and methods

### Cell culture

Mouse mammary carcinoma 4T1 cell line was purchased from Pasteur Institute of Iran cell bank. Cells were cultured in RPMI-1640 (Gibco-Invitrogene) with 10% fetal bovine serum (FBS), (Gibco-Invitrogene) and antibiotics (100U/ml penicillin and 100mg/ml streptomycin) at 37° C in a 5% CO_2_ and 95% O_2_ humidified incubator. Exhausted medium was changed every 24h.

### HBV preparation

Iranian honeybee venom was collected from *Apis mellifera* by means of an electric shocker apparatus composed of a shocker and collector unit. The shocker unit produces a light electric shock once every few seconds. Honeybees were stimulated with light electric shock and sting in beehives. The collector unit is a network of wires with small gaps and a glass plane between them. Every 25 minutes, the shocker unit turned off and the dried bee venom material on collector panel was collected by scarping.

HBV was stored as powder at −20 °C and dark condition. The main stock solution of HBV was prepared with 1mg of HBV and 1 ml phosphate buffer saline (PBS). In order to obtain a homogen and sterile solution, finally this solution was passed filter with 0.2 μm micropore. For every assay this solution was prepared freshly. For Ames assay concentration of the main stock solution was 10 mg/ml (HBV+PBS) and interested concentrations were obtained by dilution the main stock.

### Cisplatin preparation

Cisplatin was purchased from Sobhan Oncology Company of Iran. Its concentration was 50mg/ml and it was stored at 4°C and dark situation. Interested concentrations were obtained by dilution the main stock.

### MTT assay

The MTT assay is one the basis reduction of yellow MTT-dimethylthiazol diphenyl tetrazolium bromide-(tetrazole) to purple Formazan crystal by mitochondrial dehydrogenase in living cells. Adherent 4T1 cells were trypsinized by trypsin-EDTA 0.25% (Gibco-Invitrogene) then seeded in 24-well plate and cultured overnight in order to full adherence of cells to plate. The cells were treated with different concentrations of honeybee venom 0 as control, 2, 4, 6, 8, 10 μg/ml, and cisplatin, 0 as control, 5,10,15,20,25,30 μg/ml, and also cisplatin and HBV together,0 as control, 2+10, 4+10, 6+10, 8+10, 10+10 μg/ml, for 24h. Cell viability was measured using MTT assay (Sigma-America). MTT solution was prepared (5mg MTT powder in 1ml PBS) and then it was filtered. After 24h incubation of treated cells, 50μl MTT solution was added to each well, and the plate was incubated in 37°C for 4 hours and dark situation. Subsequently, the supernatant liquid was removed and 1ml DMSO, dimethylsulfoxide, (Merck, Germany) was added to each well, and plate was kept at room temperature for 15 minutes. Finally, the absorbance was measured at 570nm wavelength by a spectrophotometer (Milton Roy-Spectronic 2ID-America). The percent viability was calculated as follows:

Percent of viability = optical density of experimental group / optical density of control group x 100

### MIC assay

In order to determine the minimum inhibitory concentration of honeybee venom, the MIC assay was performed. *Salmonella* TA100 suspension in nutrient broth medium was prepared and justified by comparison against 0.5 Mc-Farland turbidity standard (1.5×10^8^ Organism/ml) tubs. The main stock solution of HBV with 10mg/ml concentration was prepared. Then the main stock was diluted and 1 mg/ml to 10 mg/ml concentrations were obtained. Finally, each tube received a certain concentration of HBV. Test tubes were incubated in 37°C for 24 h. Distilled water was used as positive control. The growth of bacteria in control and test tubes were investigated after 24h.

### Ames test

Ames test is Salmonella /microsome mutagenicity assay designed to analysis of mutagenic and anti-mutagenic factors ^14^. Salmonella TA100 used in this test have various mutations in histidine operon genes. Therefore, in the absence of histidine the bacteria are unable to grow and create colony. In presence of mutagen factors, reverse mutation is occurred so the bacteria are able to grow and form colony ^15^.

Histidine dependent strain of *Salmonella typhimurium* TA100 used for Ames test. *S. typhimurium* TA100 developed by Dr. Ames of the University of California, Berkeley, USA, was cultured in a nutrient broth (Sigma, America). The bacterial suspension was prepared 1–2 × 10^9^ cells/ml fresh culture.

In order to preparation of the rat microsomal liver enzyme (S9), mature male rats (about 200 g body weights) were deprived of food for 48h to achieve high level hepatic enzymes. Then the rats were killed and the livers were removed. After washing the livers in PBS solution, the livers were cut in to small pieces and homogenized by 1M KCl solution. Finally this solution was centrifuged for 10 min at 8700 rpm. The supernatant was isolated and stored at −80° C.

Test groups: 100μl bacteria suspension plus 100μl histidine-biotin solution (24mg biotin plus 31 mg histidine were added to 250 ml distilled water) and then 100 μl Sodium Azide solution (10ml distilled water plus 0.015 g sodium Azide) was used to the test tube contain Top Agar (0.6 g Agar plus 0.5 g NaCl plus 100ml distilled water) and finally the test tubes were incubated with 1-7 mg/ml concentrations of HBV.

Positive control: 100μl bacteria suspension plus 100μl histidine-biotin solution and then 100 μl sodium Azide solution were combined to a tube contain Top Agar.

Negative control: 100μl bacteria suspension plus 100μl histidine-biotin solution and then 100 μl distilled water were combined to a test tube contain Top Agar.

Finally, the content of these tubes after 3 second shaking were distributed on the surface of minimum medium of glucose agar (%40 glucose). The plates were incubated at 37 °C for 48 hours. All of these Anti-mutagenic assays were performed in the absence and presence of S9, and for each test three repeats were considered. Finally reversed colonies were counted and inhibition percentage was calculated by this formula:

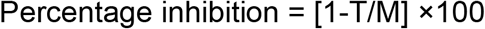

T: is the number of revertants per plate in the presence of mutagen and test sample.

M: is the number of revertants per plate in the positive control.

### Statistical Analysis

The results were assessed by one-way ANOVA method and also in combination with a Tukey test for pairwise comparison. P values less than 0.05 were considered significant. Statistical analysis was performed by SPSS 22.0 and the charts were drawn by Excel software.

## MTT Assay

For investigation of the cytotoxic effect of HBV and cisplatin on mouse mammary carcinoma 4T1 cell line, the cells were treated by various concentrations of HBV and cisplatin lonely and in combination (HBV/Cisplatin). In order to determine cell viability, MTT assay was performed. MTT assay was revealed that cisplatin and HBV have the cytotoxic effect on 4T1 cell line and they can reduce the cell viability in a dose dependent manner. As shown in figure 1.A, by increasing of HBV concentrations, the cell viability has been reduced and the treated test in comparison with control group have significant reduction of viability in the dose dependent manner (P<0.05)(figure 1.A). On the other hand cisplatin has cytotoxic effect on 4T1 cell line. High concentrations of cisplatin have shown more effective cytotoxic in comparison with the control group (P<0.001) (figure 1.B). Combination treatment of HBV and cisplatin on the 4T1 cell line showed that HBV can promote the cytotoxic effect of cisplatin in a dependent dose manner (figure 1.C). Treatment with 6μg/ml HBV and 25μg/ml with cisplatin for 24h can cause an approximately 50% 4T1 cell death. In combination cisplatin and HBV, 6μg/ml +10 μg/ml can cause approximately 50% cell death.

**Figure 1:**
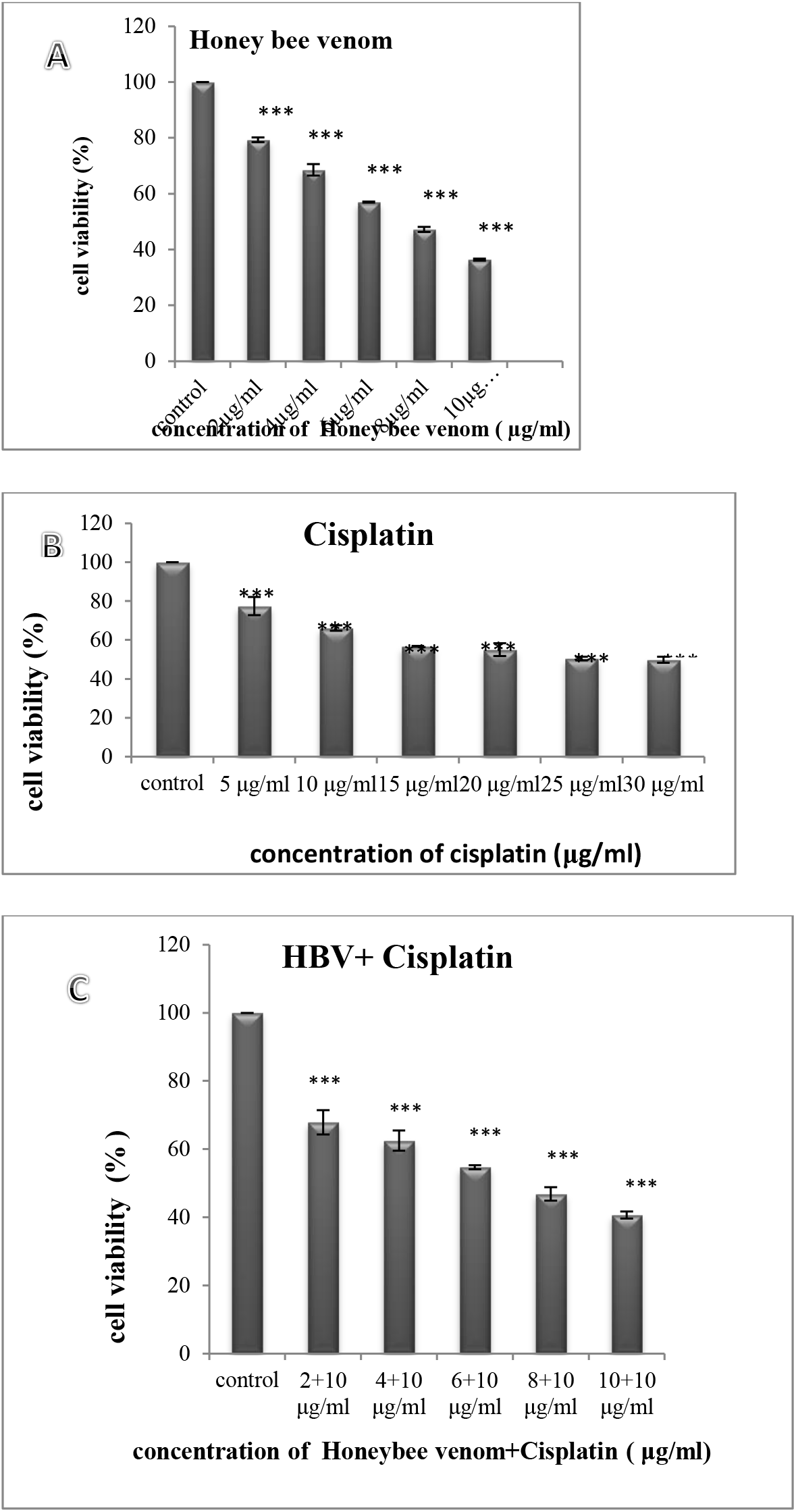
The cell viability percentages of treated 4T1 cell line with various concentrations of HBV(A), cisplatin(B) and HBV/cisplatin (C) after 24h by MTT staining (Mean ± S.E.M, *p*<0.001). HBV: Honey Bee Venom.

### MIC assay

Minimum Inhibitory Concentration assay for serial dilution concentrations of HBV was performed. Investigation of test tubes indicated that HBV can cause death in *salmonella* TA100 with dosages more than 8mg/ml. the tubes with 1mg/ml to 7mg/ml concentrations had cloudy view so the bacteria were able to grow. Hence the MIC of HBV on *salmonella* TA100 was determined 8mg/ml concentration.

### Ames assay

In order to investigate the Anti-mutagenic and also anti-cancerous activities of HBV, Ames test was performed with 1-7 mg/ml concentrations of HBV (less than MIC) in the presence and absence of S9 fraction. After 48h reversed colonies were counted (figure 2). The Plates with different concentrations of HBV have shown reduced revertants colonies in a dose dependent manner. Comparison of the test and positive control groups has shown significant differences (p<0.001) (figure 3.A). Also Ames assay was performed in the presence of S9, and result indicated that anti-mutagenic activity was improved with S9 (figure 3.B). The inhibition percentages of HBV in the presence and absence of S9 were obtained 62.76 and 56.17 (figure 4).

**Figure 2:**
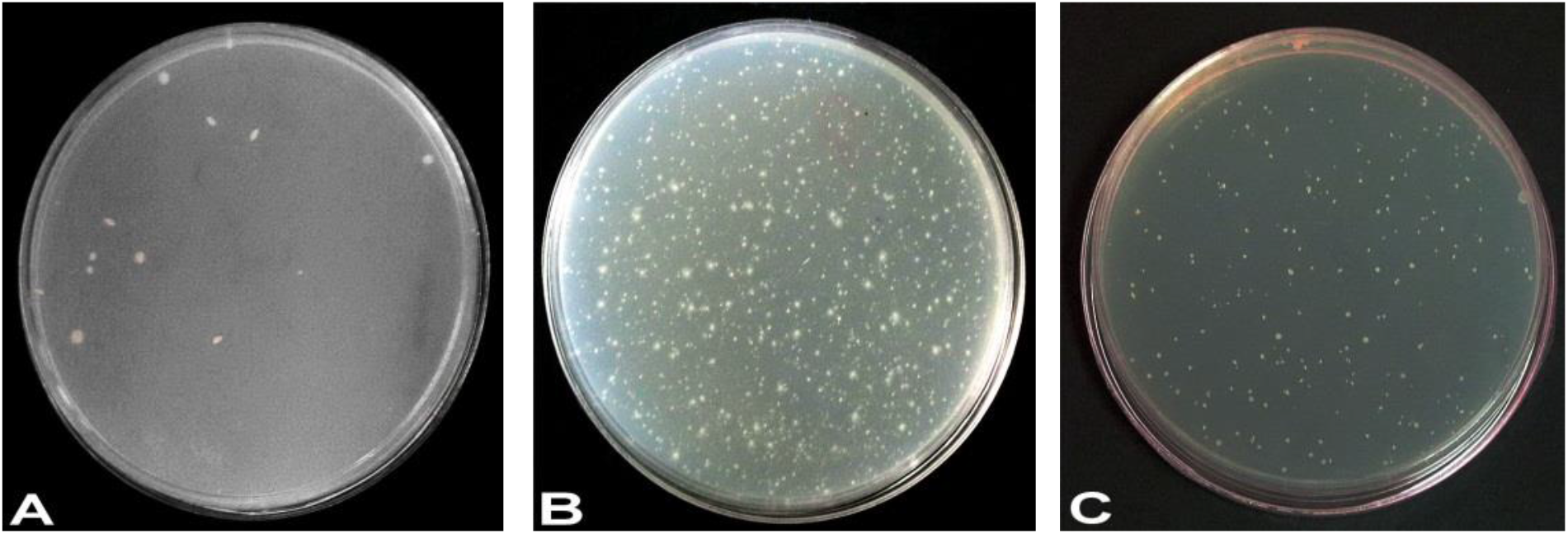
The revertants colonies in the negative test (A), positive test (B) and 7mg/ml concentration of HBV(C).

**Figure 3:**
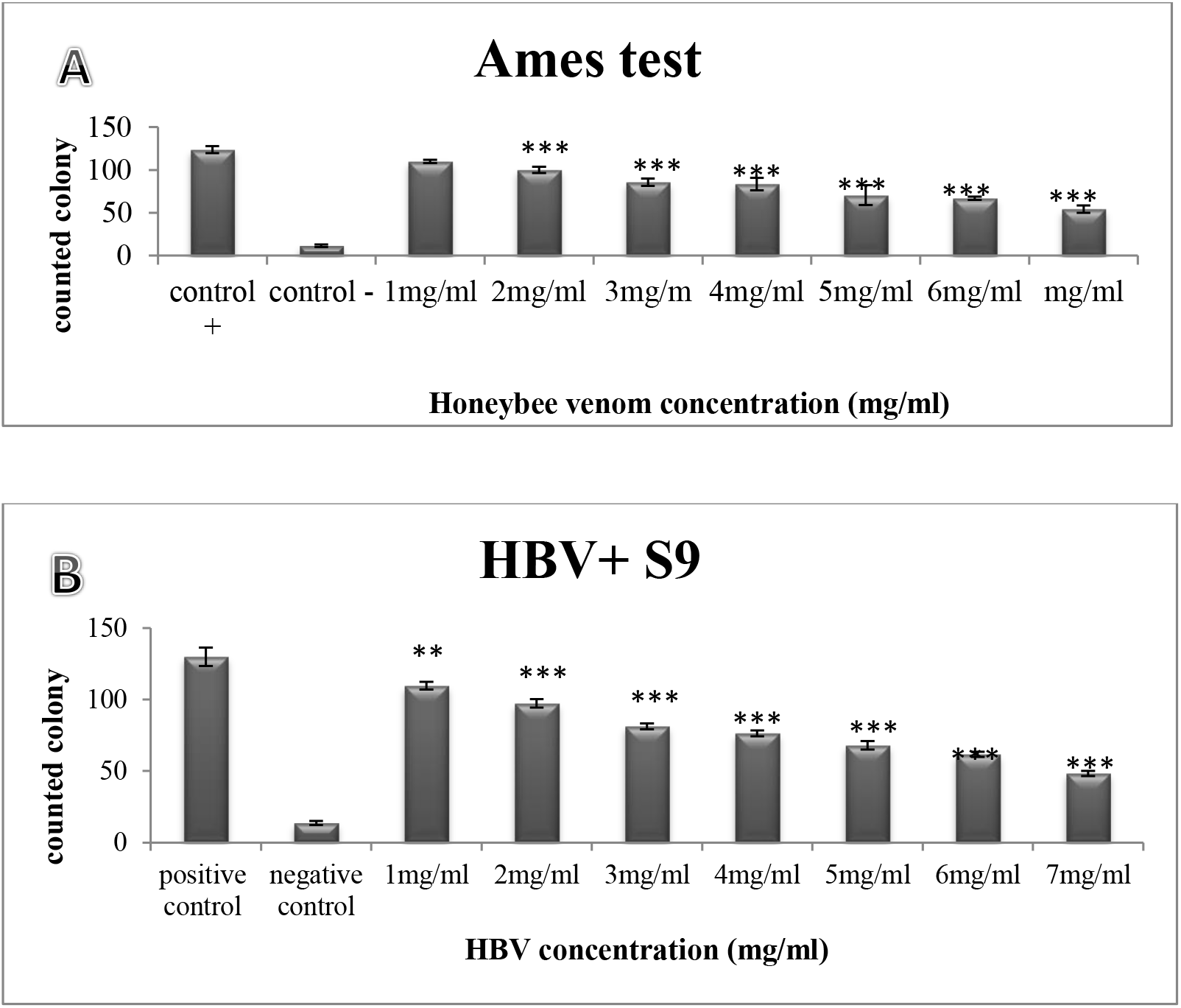
Reverted *Salmonella* TA100 colonies counts in compression with positive control group with (A) and without S9 (B) by Ames test (Mean ± S.E.M, ^***^*p*<0.001, ^**^*p*<0.01).

**Figure 4:**
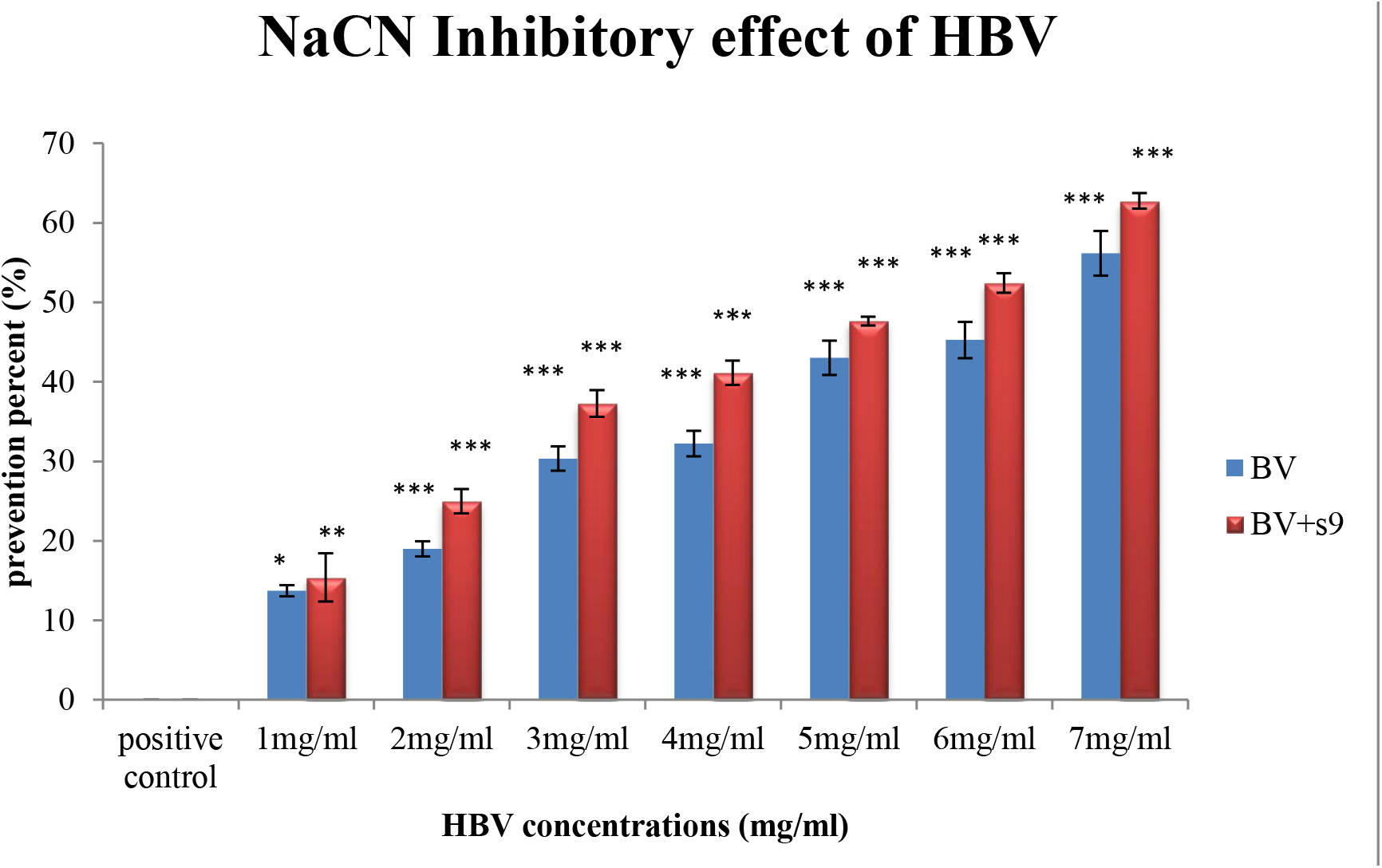
Sodium Azide inhibitory effect of HBV in comparison with the positive control group with and without S9 fraction by Ames test. S9 fraction promoted the prevention effect in a dose dependent manner (Mean ± S.E.M, ^***^*p*<0.001, ^**^*p*<0.01, ^*^*p*<0.05).

## Discussion

Breast cancer is the most prevalent cancer among women of developed countries, and its incidence has been expanding worldwide ^16^. The cytotoxic chemotherapy is used to cure the early and late stages of most cancers in the recent decade ^17^. In the 1970s, anticancer characteristic of cisplatin findings created revolution in the clinical chemotropic agents applications ^18^. In this study our objectives were examination of the cytotoxic and anti-mutagenic effects of HBV and Cisplatin, on the mouse mammary carcinoma 4T1 cell. In the present study, we report the cytotoxic activity of cisplatin on 4T1 cell line. The IC_50_ value determined for cisplatin was 25μg/ml after 24hr. The Cisplatin, cis-diamino-dichloro-platinum (II) (CDDP), a complex containing a central atom of platinum to which they have bond two ammonium ions and two chlorine ions in cis position with respect to the horizontal plane of the molecule^19^. Positive charged metals are able to bind to negative charged biomolecule. Protein and nucleic acid are excellent targets for binding to metal ions ^20^. The most of evidences suggest that the membrane proteins such as the cooper transporter 1(CTR1) accumulate Cisplatin in the cells. The replacement of water molecule with chloride ligands activates the Cisplatin molecule. The aquated forms of cisplatin can bind to DNA at the N7 position of purine bases and form primarily 1, 2-intrastrand adducts between adjacent guanosine residues ^21^. This cross linking with DNA and resulting DNA bending disrupts the replication and transcription process. Hence, the cell cycle is arrested or apoptosis is triggered ^18^. Cisplatin and other platinum complexes such as carboplatin, oxaliplatin, are more useful and effective for treating the most of cancers; however these components have several severe side effects which affect other health organs. This property results limitation of this components application ^21^. The most of new small molecules discovered in cancer treatment are Natural products or their derivatives ^22^. HBV as a natural product has had medicinal application since ancient time. HBV contains at least 18 pharmacologically active components including Melittin, Phospholipase A2, Histamine, and Adolapin. The results of various studies on the biological activity of HBV suggest that it is highly effective in several diseases such as arthritis, MS, and back pain ^11^. In our experiments the concentration of HBV that inhibited growth by 50% (IC_50_) after 24hr incubation 4T1 cell line was 6μg/ml. This data is different from the lethal dosage in other cell lines, for instances 1.43g/ml for mammary carcinoma MCa ^23^, 2μg/ml for human leukemic U937 cell line ^12^, 8μg/ml for A2780cp ^21^, 2μg/ml for human melanoma A2058 cell lines ^24^ and 10μg/ml for the human lung cancer NCI-H1299 cell line ^25^.

Hait *et al*. (2009) demonstrated that Melittin is one of the most potent inhibitors of calmodulin activity, and also a potent inhibitor of cell growth and clonogenicity. The calcium binding protein, calmodulin, plays a vital role in cellular proliferation ^26^. In 2003 Oroli investigated the effect of HBV on MCa both *in-vitro* and *in-vivo*. His results showed that HBV exerts direct (inhibition of calmodulin and prevention of cell growth) and indirect (the stimulation of macrophages and cytotoxic T lymphocytes) effect on MCa tumor cells ^27^.

Furthermore, Moon et al. (2006) studied on key regulators in HBV-induced apoptosis in human leukemic U937 cells. Their data confirmed that HBV inhibits cell proliferations in induce apoptosis in U937 cells through downregulation of Bcl-2 and upregulation of caspase-3. They demonstrated that HBV increase Fas/Fas ligand levels and decrease Cox-2 and hTERT ^12^. According to their data it can be presumed that HBV induce apoptosis through extrinsic pathway.

In 2008, Siu-Wan Ip et al. indicated that HBV induces mitochondria-dependent pathway of apoptosis in human breast cancer MCF7 cells. Their results confirmed that HBV induces DNA strand breaks and promoted P53 and P21 factors. HBV also affects the ratio of Bax/Bcl-2 level leading to releasing cytochrome c and finally triggering of mitochondrial apoptosis pathway ^28^.

Accordingly, our findings are in agreement with other reports about the cytotoxic effect of HBV on cancerous cells. Since HBV probably targets the DNA molecule and also inhibit calmodulin protein it is probable that HBV exerts its cytotoxic and growth prevention effects in 4T1 cell line through intrinsic/ extrinsic apoptosis pathway or cell cycle arrest.

Our findings revealed that cytotoxic effect of Cisplatin are potentiated by combination of non-lethal concentrations of HBV. In 2009 Orsolic investigated the cytotoxic effect of BV in combination with Bleomycin in Hela and V79 cell lines. The non-lethal dosage of HBV with bleomycin cause increase death cell in dose-depended manner. He presumed that HBV inhibits DNA repair and this may be the mechanism by which it increases bleomycin lethality and inhibits recovery from bleomycin-induced damage ^23^. On the other hand, Gajski (2008) demonstrated that HBV induces single- and double-strand DNA breaks in human lymphocytes ^29^. According to these findings and since cisplatin targets the DNA molecule it is probable that HBV is able to promote the cytotoxic effect of cisplatin via this mechanism. Our data in this study are in direction with Alizadeh et al. study on investigation of the synergistic effect of BV and cisplatin on human ovarian cancer cell line A2780cp in which HBV potentiated the cytotoxic effect of cisplatin ^21^.

Mutation, a natural process that changes a DNA sequence, is one of the most important causes of cancer. Also changes in the structure of chromosomes have critical role to create the most of malignancies ^31^. In this study HBV inhibited the reverted mutation in dose dependent manner. According to Ames theory the number of revertants colony in positive control plate (with carcinogen substrate) should be 2 times more than test plates. Also mutagen prevention percentage ranges have 3 classes interpretation include inhibitory percent more than 40% which show high prevention potential, between 25-40% which show medium potential and less than 25% which show negative prevention ^31^. Hence, in this study, our results showed that the number of revertants colony in the test plate with 7 mg/ml concentration of HBV (with S9 and without S9) is equal to less than half of the number of colony in Positive control plate. On the other hand the inhibitory percent for 5mg/ml, 6 mg/ml and 7 mg/ml concentrations of HBV Respectively is 43.05%, 45.26%, and 56.17% in absence of S9, so these HBV concentrations showed high prevention potential. Furthermore, the S9 fraction potentiated HBV inhibitory effect. The S9 fraction is prepared from the liver of rats and it contains hepatic enzyme which effectively bioactive promutagens to mutagens. This metabolic activation of mutagens is considered vital step for carcinogenesis, because most of carcinogen must enzymatically transformed to electrophilic specious. This activated mutagen is able to covalently bind to DNA molecule leading to mutation ^32, 33^ Other studies were performed to investigate antimutagenic some natural products. In 2011 Ghazali *et al*. confirmed that extract of *M. speciosa* indicates anti-mutagenic activity ^34^ Also Issazadeh and his coworkers (2012) concluded that olive leaf shows anti-mutagenic and anti-carcinogenic effects ^30^. Also our data demonstrated that in the presence of S9 fraction, the mutagen prevention percent of HBV was promoted. Since S9 fraction activate sodium azide mutagen so it is likely that HBV prevents the mutagenic effects of sodium azide. Finally, our result in agreement with previous studies and its interpretation confirmed that HBV has anti-mutagenic and anti-cancerous effect in dependent dose manner.

### Conclusion

According to our results in this study, we assessed that HBV as a natural product has cytotoxic effect on mouse mammary carcinoma 4T1 cell line. Cisplatin as a chemotropic drug to cure several types of cancer has cytotoxic effect on 4T1 cell line, but in combination with HBV, its cytotoxic effect is potentiated and it can be more effective in non-lethal dosages. Furthermore, anti-mutagenic and anti-cancerous activities of HBV were seen in the presence of S9 metabolic activation system in the all concentrations of HBV. It will be an excellent perspective to innovate approaches to prevent and treat the some features of cancer.

## Acknowledgment

This project was performed in the Laboratory of Cell and Developmental Biology at Kharazmi University and the authors would like to thank all lab members, Masoumezaman Alizadehnohi, Siamak Yari, of for their supports.

